# Evaluating the power and limitations of genome-wide association mapping in *C. elegans*

**DOI:** 10.1101/2021.09.09.459688

**Authors:** Samuel J. Widmayer, Kathryn Evans, Stefan Zdraljevic, Erik C. Andersen

## Abstract

A central goal of evolutionary genetics in *Caenorhabditis elegans* is to understand the genetic basis of traits that contribute to adaptation and fitness. Genome-wide association (GWA) mappings scan the genome for individual genetic variants that are significantly correlated with phenotypic variation in a population, or quantitative trait loci (QTL). GWA mappings are a popular choice for quantitative genetic analyses because the QTL that are discovered segregate in natural populations. Despite numerous successful mapping experiments, the empirical performance of GWA mappings has not, to date, been formally evaluated for this species. We developed an open-source GWA mapping pipeline called NemaScan and used a simulation-based approach to provide benchmarks of mapping performance among wild *C. elegans* strains. Simulated trait heritability and complexity determined the spectrum of QTL detected by GWA mappings. Power to detect smaller-effect QTL increased with the number of strains sampled from the *C. elegans* Natural Diversity Resource (CeNDR). Population structure was a major driver of variation in GWA mapping performance, with populations shaped by recent selection exhibiting significantly lower false discovery rates than populations composed of more divergent strains. We also recapitulated previous GWA mappings of experimentally validated quantitative trait variants. Our simulation-based evaluation of GWA performance provides the community with critical context for pursuing quantitative genetic studies using CeNDR to elucidate the genetic basis of complex traits in *C. elegans* natural populations.

## INTRODUCTION

Quantitative trait variation in human populations is abundant and arises from genetic differences between individuals, as well as complementary or detrimental inputs from the environment. Genetic variation can be statistically linked to phenotypic variance using genome-wide association studies (GWAS). GWAS have uncovered genetic variants that contribute cumulatively to human disease risk and complex trait variation (Visscher *et al*. 2017). However, the most powerful and useful applications of GWAS to complex human traits rely on precise phenotype measurements from hundreds of thousands of individuals. The subsequent statistical penalties for multiple comparisons increase as the scale of GWAS increases. Also, many important sources of variation in disease risk and trait variation cannot be measured ethically, reliably, and with sufficient statistical power in human populations (*e.g.*, cellular pathology underlying behavioral traits and variation in diet or xenobiotic exposure underlying metabolic traits). Finally, GWAS studies have a historical underrepresentation among non-White ethnic groups created in part by healthcare inequities, which cause polygenic risk scores among these groups to be significantly less accurate (Martin *et al*. 2019). This gap underscores an urgent need for replicable and translatable GWA platforms with the added ability to dissect traits that are difficult to assay in humans.

The development of genetic reference populations of several organisms has become increasingly popular and has facilitated the analysis of complex traits. Notable examples of this include the *Drosophila* Synthetic Population Resource (King *et al*. 2012a; b)*, Drosophila* Genetic Reference Panel (Mackay *et al*. 2012), the Collaborative Cross (Churchill *et al*. 2004; Chesler *et al*. 2008; Aylor *et al*. 2011) and Diversity Outbred (Svenson *et al*. 2012; Churchill *et al*. 2012) mouse populations, the hybrid mouse diversity panel for association mapping (Bennett *et al*. 2010), *Arabidopsis* MAGIC and recombinant inbred lines (Kover *et al*. 2009; Klasen *et al*. 2012), and nested association mapping lines in both maize (Yu *et al*. 2008; McMullen *et al*. 2009) and sorghum (Bouchet *et al*. 2017). These genetic reference populations offer tremendous benefits for quantitative genetics because they take advantage of well characterized genomic resources, repeated measurements that can be collected from multiple genetic backgrounds, and population-wide measurements across diverse individuals that can be made in controlled environments. The free-living roundworm nematode *Caenorhabditis elegans* has contributed to discoveries at every level of biology, has rich genomic resources, and can be easily genetically manipulated. Over the past few decades, the number of catalogued genetically unique *C. elegans* isolates has expanded, giving rise to diverse collections of strains useful for quantitative genetics (Cook *et al*. 2017; Lee *et al*. 2021). For example, the *C. elegans* Multiparent Experimental Evolution (CeMEE) lines offer fertile ground for quantitative trait locus (QTL) mapping with high-resolution and detection power (Noble *et al*. 2017, 2021). Although rich in novel haplotypes, the CeMEE panel represents only a fraction of the genetic variation present across the *C. elegans* species. Separately, since the generation of the CeMEE panel, the *C. elegans* Natural Diversity Resource (CeNDR) has expanded to over 500 unique *C. elegans* strains. Genome-wide association (GWA) mapping has repeatedly linked phenotypic variation of all types to alleles segregating among these strains (Ghosh *et al*. 2012; Ashe *et al*. 2013; Cook *et al*. 2016; Zdraljevic *et al*. 2017, 2019; Lee *et al*. 2017, 2019; Laricchia *et al*. 2017; Hahnel *et al*. 2018; Webster *et al*. 2019; Gimond *et al*. 2019; Na *et al*. 2020; Evans *et al*. 2020, 2021a; b; Zhang *et al*. 2021). However, GWA mapping has not, to date, been formally evaluated for its power and precision to detect QTL across a range of genetic architectures.

The ability to identify functional natural variation in complex traits in *C. elegans* using genome-wide association is confounded by idiosyncratic genomic features. For instance, adaptation to human-associated habitats is hypothesized to have caused the generation of haplotypes with signatures of selective sweeps among many wild *C. elegans* strains. Within these swept haplotypes, genetic variation is drastically reduced and long-range linkage disequilibrium is high - sometimes stretching over 85% of whole chromosomes (Andersen *et al*. 2012). Approximately 66% of the *C. elegans* strains available in CeNDR contain at least one chromosome of which at least 30% can be categorized as a swept haplotype. The unintended consequence in GWA mapping is that, if the phenotype of interest happens to segregate with a common swept haplotype, it is likely that insufficient ancestral recombination has occurred across the associated swept haplotype to resolve single candidate loci. By contrast, *C. elegans* strains from Hawaii harbor nearly three times the levels of genetic diversity of non-Hawaiian strains and often lack signatures of recent selection in spite of recent migration and gene flow (Crombie *et al*. 2019). Furthermore, genetically distinct *C. elegans* strains contain “hyperdivergent” regions (Thompson *et al*. 2015) (regions of the genome characterized by high allelic diversity and, therefore, uncertainty in gene content compared to the N2 reference genome) that segregate at varying frequencies. These regions are hypothesized to be maintained by balancing selection and are predicted to harbor alleles for biological processes that are crucial for environmental sensing, pathogen responses, and xenobiotic stress responses (Lee *et al*. 2021). These observations suggest that evolutionary biology is inextricable from GWA mapping performance in *C. elegans* and that the conclusions drawn about complex trait variation from these analyses are dictated by the population structure of the mapping population. However, the magnitude of the effect of population structure and segregating hyperdivergent regions on mapping performance has not been quantified. In order to assess how mapping performance varies as a function of population composition, we require an approach that can rapidly simulate GWA mappings and address important caveats unique to *C. elegans* genome biology.

We have developed NemaScan, an open-source pipeline for GWA mapping in *C. elegans*. NemaScan offers two profiles: a mapping profile where users can supply population-specific variant information and a phenotype to perform their own analyses on real data and a simulation profile where users can supply a variety of parameters to provide baseline performance benchmarks for a past, present, or prospective experiment. These parameters include trait heritability, polygenicity, a minimum minor allele frequency for variants included in the marker set, custom sample populations, and specific regions of interest where QTL are simulated and mapped iteratively. NemaScan makes use of two different formulations of the genomic relationship matrix in attempts to correct for varying types of population structure known to exist across the *C. elegans* species. We present empirical estimates of detection power and false discovery rates derived from the simulation profile for GWA mapping across different genetic architectures, and we confirm that GWA mappings in *C. elegans* robustly identify most large-effect QTL. We also demonstrate that GWA performance in *C. elegans* is improved by both increasing the number of strains tested in a population and homogenizing the genetic makeup of the population in question with respect to swept haplotypes. Finally, we quantify the precision of GWA mapping when QTL are present on different chromosomes and within hyperdivergent regions that segregate in swept and divergent populations. These performance benchmarks provide the *C. elegans* community with critical context for interpreting the results of ongoing quantitative genetic studies using CeNDR, and in so doing, increase our understanding of the genetic basis of complex traits in *C. elegans*.

## MATERIALS AND METHODS

### Additions to the Caenorhabditis elegans Natural Diversity Resource (CeNDR)

CeNDR is composed of 1,379 unique *C. elegans* isolates. The process of isolating and identifying unique *C. elegans* strains, generating whole-genome sequence data, and calling high-quality variants has been described in-depth previously (Crombie *et al*. 2019; Lee *et al*. 2021). Briefly, nematodes that could be unambiguously described as *C. elegans* by both morphological characteristics and ITS2 sequencing were reared, and genomic DNA from these strains (*n =* 1238) was isolated and whole-genome sequenced. High-quality, adapter-trimmed sequencing reads were aligned to the N2 reference genome and SNVs were called for each strain using BCFtools. After variant quality filtering, the pairwise genetic similarity of all strains is considered. Strains which share alleles across at least 99.97% of all segregating sites are considered members of the same isotype group. After measuring concordance among all strains, 540 unique isotype groups were identified. In this manuscript, we use the term “strain” to refer to each strain chosen to represent the collection of genetically similar strains within that isotype group (*i.e.*, “isotype reference strain”). All data used in GWA mapping simulations (isotype-level hard-filtered SNVs, sweep haplotype calls, and hyperdivergent region calls) were downloaded from the 20210121 CeNDR release (https://www.elegansvariation.org/data/release/latest).

### Genome-wide association (GWA) mapping simulations

All GWA mapping simulations were completed using the simulation profile of the NemaScan pipeline, available at https://github.com/AndersenLab/NemaScan. The VCF file was then pruned for variants in *r^2^* ≥ 0.8 within 50 kb windows obtained in ten-variant steps and filtered to contain variants with a minor allele frequency greater than or equal to the user-supplied minor allele frequency cutoff. The LD-pruned and MAF-filtered VCF was then used to construct a genomic relationship (kinship) matrix among all strains using the --make-grm and -- make-grm-inbred function from GCTA. The algorithm for constructing the genomic relationship matrix and its benefits for association mapping has been described in-depth elsewhere (Jiang *et al*. 2019). Separately, the user-specified number of causal variants are then sampled from LD-pruned and MAF-filtered VCF and assigned effects sampled from the user-specified effect distribution (either *Uniform [a,b]* (where a = the user-specified minimum effect and b = the user-specified maximum effect) or *Gamma* (*k =* 0.4, *θ* = 1.66)). Once these effects were assigned to causal variants, phenotype values were then simulated for each of the strains in the supplied population using the --simu-causal-loci function from GCTA and the user-specified trait heritability. Simulated phenotypes, filtered variants, and the genomic relationship matrix were brought together to perform rapid GWA using the --mlma-loco and --fastGWA-lmm-exact functions by GCTA. The former function accepts a limited sparse kinship matrix composed of all chromosomes except the chromosome containing the tested marker (LOCO = “leave one chromosome out”), and the latter accepts a full sparse kinship matrix specifically calculated for inbred model organisms.

### Performance Assessment

Raw mapping results were aggregated by finding the lowest *p-*value for each marker comparing the GWA mapping results from both functions. This aggregation step is performed to take advantage of the benefits provided by the LOCO approach and the inbred kinship matrix simultaneously. The aggregated mapping results were then processed to determine whether each SNV exceeds the user-specified threshold of statistical significance. The user has three choices of significance thresholds: i) Bonferroni correction using all tested markers (“BF”), ii) Bonferroni correction using the number of independent tests determined by eigendecomposition of the population VCF (“EIGEN”), or iii) any nominal value supplied by the user. The phenotypic variance explained by each SNP was also calculated using a simple ANOVA model using the simulated phenotypes as a response and the allelic state of each strain as a factor. SNVs exceeding the user-specified significance threshold were then grouped into QTL “regions of interest”, motivated by the fact that *C. elegans* can be rapidly crossed to generate NILs harboring small introgressed regions to localize candidates using fine mapping. Regions of interest were determined by finding significantly associated markers within one kilobase of one another. Once no more markers met this criterion, the region of interest was extended on each flank by a user-specified number of markers. The QTL region of interest was denoted by the peak association found within the region and was assigned the phenotypic variance explained by that peak marker and its frequency in subsequent analyses.

We then cross-referenced simulated causal variants for each mapping and asked whether any detected QTL region of interest overlapped with a simulated causal variant. The possible outcomes regarding the performance of GWA mapping to detected simulated causal variants were (1) a simulated causal variant was significantly associated with phenotypic variation and was the peak association within a region of interest, (2) a simulated causal variant was significantly associated with phenotypic variation but was *not* the peak association within a region of interest, (3) a simulated causal variant was *not* significantly associated with phenotypic variation but still fell within a QTL region of interest, and (4) a simulated causal variant was neither associated with phenotypic variation nor fell within a QTL region of interest. For each replicate mapping, we calculated detection power as the number of causal variants that adhered to criteria (1) or (2) and divided them by the total number of causal variants simulated for that mapping. QTL regions of interest that did not contain a simulated causal variant were tabulated as false discoveries, and the false discovery rate (FDR) was calculated as the number of QTL regions of interest that did not contain a simulated variant divided by the total number of QTL regions of interest for each mapping. For analyses assessing the ability of GWA mappings to detect causal variants explaining a particular amount of phenotypic variance, detection power was calculated by first determining the number of causal variants that adhered to criteria (1) or (2) *and* that explained that amount of phenotypic variance. We then divided them by the total number of causal variants simulated that explained the same amount of phenotypic variance across *all* mappings (instead of individual replicates).

### Demographic Characterization of Strains

Haplotype data for 540 *C. elegans* strains was obtained from the 20210121 CeNDR release. The degree of swept haplotype sharing among strains was determined in a similar fashion to that previously described (Crombie *et al*. 2019; Lee *et al*. 2021; Zhang *et al*. 2021). Briefly, the length of every haplotype present in each strain was recorded, and if regions sharing the most common haplotype were longer than 1 Mb, these haplotypes were recorded as swept haplotypes. Haplotypes outside of these highly shared regions were recorded as divergent haplotypes. Only swept haplotypes on chromosomes I, IV, V, and X were considered in strain classification because selective sweeps are not found on chromosomes II and III. If swept haplotypes composed greater than or equal to 30% of the length of these chromosomes, that chromosome was considered swept. Swept strains were determined as those strains that contain at least one swept chromosome, and divergent strains are those strains that do not. In total, 357 swept and 183 divergent strains were identified. Some populations used in simulations were constructed by sampling among these swept and divergent strains (**Figure 3**), and others were sampled from the overall collection of 540 strains (**Figure 2, Figure 3**). In simulations comparing QTL simulated in hyperdivergent regions from those simulated outside of such regions, we compared 182 swept strains to 183 divergent strains selected on the basis of containing at least 37 hyperdivergent regions, regardless of their population frequency. Dendrograms representing population differentiation were constructed for these swept and divergent populations by filtering genetic variants identically to NemaScan and passing these variant calls to vcf2phylip (Ortiz 2019) and QuickTree (https://github.com/khowe/quicktree).

### Statistical Testing

Determinations of significant differences in performance among experimental factors were determined using both parametric and non-parametric specifications of power or empirical FDR as a response. Simulation regimes where only one QTL was specified for each simulated mapping resulted in a binary distribution of power output, and therefore differences in performance as a function of experimental factors were determined using the Kruskall-Wallis test. Differences between all pairwise contrasts of factor levels were determined using the Dunn’s test. In cases where multiple experimental factors were considered simultaneously (for example, whether mapping strain set and the location of the single simulated QTL *interacted* to determine performance), factors were combined to make an aggregate factor and tested using the Kruskall-Wallis test. When the specified number of QTL were greater than one, differences in performance as a function of single and multiple factors were determined using the One-Way ANOVA and Two-Way ANOVA tests, respectively, and followed up with *post hoc* tests using Tukey’s HSD.

### Data Availability

The simulation and mapping profiles of NemaScan are available for download at https://github.com/AndersenLab/NemaScan and are accessible with the same pipeline. Users are invited to use NemaScan to perform GWA mappings on their own traits of interest or leverage the simulation framework to explore the potential of GWA for their own traits of interest or to assess the likelihood of previous mapping results. In addition, all parameter specifications used to generate the mappings in this manuscript are contained in **Supplemental Table 1**. All code and data used to replicate the data analysis and figures presented are available for download at https://github.com/AndersenLab/nemascan_manuscript. All variant calls, hyperdivergent region calls, and selective sweep haplotype calls are available at https://www.elegansvariation.org/data/release/latest. Finally, prospective users are also encouraged to use NemaScan to perform their own mappings at https://www.elegansvariation.org/mapping/perform-mapping/.

## RESULTS

### GCTA software improves C. elegans GWA power and precision

The previous GWA mapping workflow, cegwas2-nf (Zdraljevic *et al*. 2019), was built on the foundation of kinship matrix specification using EMMA or EMMAX (Kang *et al*. 2008, 2010) implemented by R/rrBLUP (Endelman 2011) as the association mapping algorithm. However, with the advent of more efficient and flexible algorithms, we wondered whether GCTA offered better performance. We first optimized the algorithm used for fitting linear mixed models and estimating kinship among individuals in the GWA mapping. Simulations were performed using four different association mapping algorithms, of which three are different implementations of association mapping using GCTA software (Yang *et al*. 2011; Jiang *et al*. 2019). (1) EMMA: GWA mapping using R/rrBLUP fits a kinship matrix and performs association using variance components using the “P3D = TRUE” option. (2) LMM-EXACT-LOCO: GCTA-LOCO fits a kinship matrix constructed using all chromosomes except for the chromosome harboring the tested genetic variant (“leave one chromosome out”). (3) LMM-EXACT: fastGWA fits with a sparse kinship matrix using all chromosomes. (4) LMM-EXACT-INBRED: fastGWA fits a sparse kinship matrix tailored towards populations composed of inbred organisms.

We next used convenient features offered by GCTA to simulate quantitative traits (-- simu-qt) and assign effects to QTL (--simu-causal-loci) across a panel of real *C. elegans* genomes. The statistical properties of each mapping algorithm have been reported elsewhere (Yang *et al*. 2011; Jiang *et al*. 2019). To begin, we used a population of 203 isolates that were previously measured for susceptibility to albendazole (Hahnel *et al*. 2018). We simulated 50 quantitative traits with increasing narrow-sense heritability (the proportion of phenotypic variance explained by specific genetic differences between strains, *h^2^*), ranging from 0.1 to 0.9, supported by either a single QTL or five independent QTL. Each QTL was assigned a large effect size sampled from a uniform distribution (**Supplemental Figure 1**) to increase the likelihood that at least one true QTL was detected in each simulation.

We measured the statistical power and the empirical false discovery rate (FDR; the proportion of detected QTL regions that lack a simulated causal variant exceeding the multiple testing correction significance threshold) of each association mapping workflow across varying levels of trait heritability and for traits supported by either one or five QTL. We observed that GCTA-based workflows were more powerful than EMMA for almost every simulated genetic architecture (**Supplemental Figure 2A**). When mapping a single causal QTL, we observed that algorithms exhibited almost identical power when that QTL explained at least 30% of the phenotypic variance (Kruskall-Wallis test, *p* ≥ 0.295). However, when traits were supported by five QTL, power varied among algorithms and increased as a function of trait heritability. When *h^2^* < 0.4, the algorithms exhibited no significant differences in detection power (Kruskall-Wallis test, *p* ≥ 0.276). When *h^2^* ≥ 0.4, algorithms diverged in performance, with LMM-EXACT and LMM-EXACT-INBRED algorithms generally exhibited lower power than both the EMMA and LMM-EXACT-LOCO algorithms (Dunn test, *p_adj_* ≤ 0.01385). Furthemore, the LMM-EXACT-LOCO algorithm exhibited significantly greater power than EMMA for traits with *h^2^* > 0.7 (Dunn test, *p_adj_* ≤ 0.00826) (**Supplemental Table 2**). We also observed only modest differences in empirical false discovery rates (FDR) among algorithms at different trait heritabilities, among them being that the LMM-EXACT-LOCO and LMM-EXACT-INBRED algorithms often exhibited lower empirical FDR than both the EMMA and LMM-EXACT algorithms (**Supplemental Figure 2B**, **Supplemental Table 3**). These results indicated that mapping algorithms implemented by GCTA have equal or greater power for QTL detection and lower FDR in *C. elegans* than the previous implementation of GWA mapping using EMMA.

The observation that either the LMM-EXACT-LOCO or LMM-EXACT-INBRED algorithms exceeded the QTL detection power of EMMA across a range of trait heritabilities motivated us to integrate both mapping algorithms into new simulation and mapping profiles. In future simulations presented here and in the mapping workflow available on CeNDR, traits are mapped using both the LMM-EXACT-LOCO and LMM-EXACT-INBRED algorithms, and mapping results from each are combined by taking the lower *p*-value from each algorithm’s association test for every marker. Although this approach may inflate the FDR for a given mapping, we prioritized a more flexible range of detection power in order to provide researchers with greater potential for QTL discovery for diverse types of traits and differentially stratified populations given that the algorithms specify genetic covariance differently. Mapping results provided using CeNDR include the combined mapping results with metadata, as well as raw individual mapping outputs for both algorithms if researchers prefer the handling of the genomic relatedness from one algorithm over the other. This combined output integrated into distinct simulation and mapping profiles is the foundation of our new GWA mapping workflow, called NemaScan.

### Genetic architecture dictates the spectrum of C. elegans QTL detection using GWA mapping

One of the most critical benchmarks for GWA mapping in *C. elegans* is the number of QTL underlying complex traits that can be detected. Traits of particular interest are noisy or highly sensitive to environmental perturbations, controlled by many genes with relatively small effects, or controlled by collections of alleles at varying frequencies in the sample population. In order to quantify the ability of NemaScan to identify QTL in natural populations of wild isolates, we performed simulations making changes to the genetic architectures of simulated traits. First, simulated QTL effects were drawn from a *Gamma* (*k =* 0.4, *θ* = 1.66) distribution, conforming to the assumption that the natural genetic variants underlying complex traits and adaptation primarily contribute small phenotypic effects but occasionally exert moderate or large effects (Supplementary Figure 3). Second, because experimenters have limited control over the realized heritability of their trait of interest, traits were simulated with *h^2^* = 0.2, 0.4, 0.6, or 0.8. For each heritability specification, traits were either supported by 1, 5, 10, 25, or 50 QTL to examine GWA performance across a broad spectrum of genetic architectures. Third, we simulated each of these genetic architectures in the complete set of 540 wild isolates currently available from CeNDR to determine the expected performance in the theoretical case where every available genetic background is assayed for a phenotype of interest.

We observed that detection power decreased as a function of the number of supporting QTL for each simulated trait, regardless of its heritability. In the simplest case where a single QTL accounted for all of the phenotypic variance, mappings exhibited at least 97% power to detect it on average. However, detection power decreased as simulated trait complexity increased, especially for less heritable traits (**Figure 2A**). NemaScan exhibited only 33.2% power to detect five QTL architectures and only 7.6% power to detect 50 QTL architectures, corresponding to detecting on average 1.66 true QTL out of five or 3.78 true QTL out of 50, respectively. Depending on the number of simulated QTL, detection power increased by between a two-fold (five QTL) to six-fold (50 QTL) magnitude by increasing trait heritability from 0.2 to 0.8. The empirical FDR also decreased as a function of genetic complexity (**Figure 2B**).

Mappings of five QTL architectures produced a mean FDR of 11.8%, and mappings of 50 QTL architectures produced a mean FDR of 0.41%. Among traits supported by the same number of QTL, FDR increased with trait heritability but to a much lesser extent than detection power. These results demonstrated that features of complex traits that alter performance of GWA mappings in other model systems generally also extend to relatively small *C. elegans* sample populations. By quantifying increases in power and FDR across various genetic architectures, we also provide performance benchmarks for GWA mappings in *C. elegans* and emphasize that obtaining more precise phenotype measurements, and thereby reducing environmental noise, improves the prospects of precise QTL detection across *C. elegans* strains.

In *C. elegans* as well as other systems, the power to detect causal alleles underlying QTL in natural populations is limited in part by their frequency and effect size, which together contribute to the fraction of phenotypic variance explained by that QTL. We calculated the phenotypic variance explained by each causal QTL across all simulations and found that true positive QTL (simulated QTL with significant trait associations) had significantly greater explanatory power than false negative QTL (causal QTL without significant trait associations) within all combinations of trait heritability and polygenicity regimes (One-Way ANOVA, Tukey HSD, *p_adj_* < 0.05) except for one QTL and *h^2^* = 0.2 (One-Way ANOVA, Tukey HSD, *p_adj_* ≥ 0.962) (**Figure 2C**). We also observed that the simulated variance explained by significantly associated true positive markers was significantly different among all trait heritability and polygenicity combinations. The median simulated variance explained by top hits in polygenic architecture simulations ranged from 7.41% (*h^2^* = 0.2; 50 QTL) to 42.35% (*h^2^* = 0.8; five QTL), and the median simulated variance explained by false negative QTL consistently remained below 2%. When markers with the highest statistical association were also the causal markers, they explained significantly more phenotypic variance than significantly associated causal markers that were not peak associations (One-Way ANOVA, Tukey HSD, *p_adj_* < 0.05), except for traits supported by one QTL (One-Way ANOVA, Tukey HSD, *p_adj_* ≥ 0.073). We conclude from these patterns that QTL detected through GWA mapping in *C. elegans* were indeed enriched for alleles with outsized effects on trait variation, explaining smaller amounts of the total trait heritability as trait complexity increased.

### Sample size and population structure modulates the sensitivity of GWA mapping in C. elegans

A common practical limitation of the scope and performance of any GWAS is the size of the sample population for which phenotypes have been measured. *C. elegans* GWA mappings are no exception, despite high-throughput phenotypic platforms becoming more commonplace in studies of natural phenotypic variation (Yemini *et al*. 2013; Andersen *et al*. 2015). We quantified the detection power of NemaScan when applied to complex traits given the finite sampling potential of a typical GWA experiment. To accomplish this simulation, we subsampled the 540 CeNDR isolates at five different depths (*n =* 100, 200, 300, 400, or 500) 50 times each. We then measured the sensitivity of GWA mappings to detect simulated QTL according to the phenotypic variance that they explained by grouping simulated QTL into bins representing increasing influence on trait variation. Among all QTL simulated, we found no clear differences in minor allele frequencies among populations of different sizes (**Supplemental Figure 4**).

We first observed that, as expected, overall detection power generally increased as a function of sampling depth. The average power to detect five QTL among 100 subsampled strain mappings was 0.33 ± 0.15 (roughly one QTL out of five), increasing to 0.46 ± 0.18 (at least two QTL out of five) among 500 subsampled strain mappings (**Table 1**). The observation of roughly 46% power to detect five QTL at *h^2^* = 0.8 among 500 subsampled strains is consistent with our previous simulation results (**Figure 1A**) and indicates that as the number of strains in CeNDR expands so will the potential for NemaScan to detect all of the QTL for a given trait. We also observed that the impact of increasing sample size was most striking when considering the sensitivities of mappings to detect QTL with smaller effects (**Figure 2**). Both 100-strain and 500-strain mappings had greater than 80% power to detect QTL that explained greater than 50% of the phenotypic variance. However, the power of 500-strain mappings to detect QTL explaining as little as 7.5% of the phenotypic variance (0.52 ± 0.2) was nearly five times greater than that of 100-strain mappings (0.11 ± 0.1) (**Supplemental Table 4**). These results indicate that power to detect QTL with large effects increased only marginally with increasing sampling depth, and power to detect QTL with smaller effects improves significantly by adding more strains to mapping populations.

**Figure 1:**
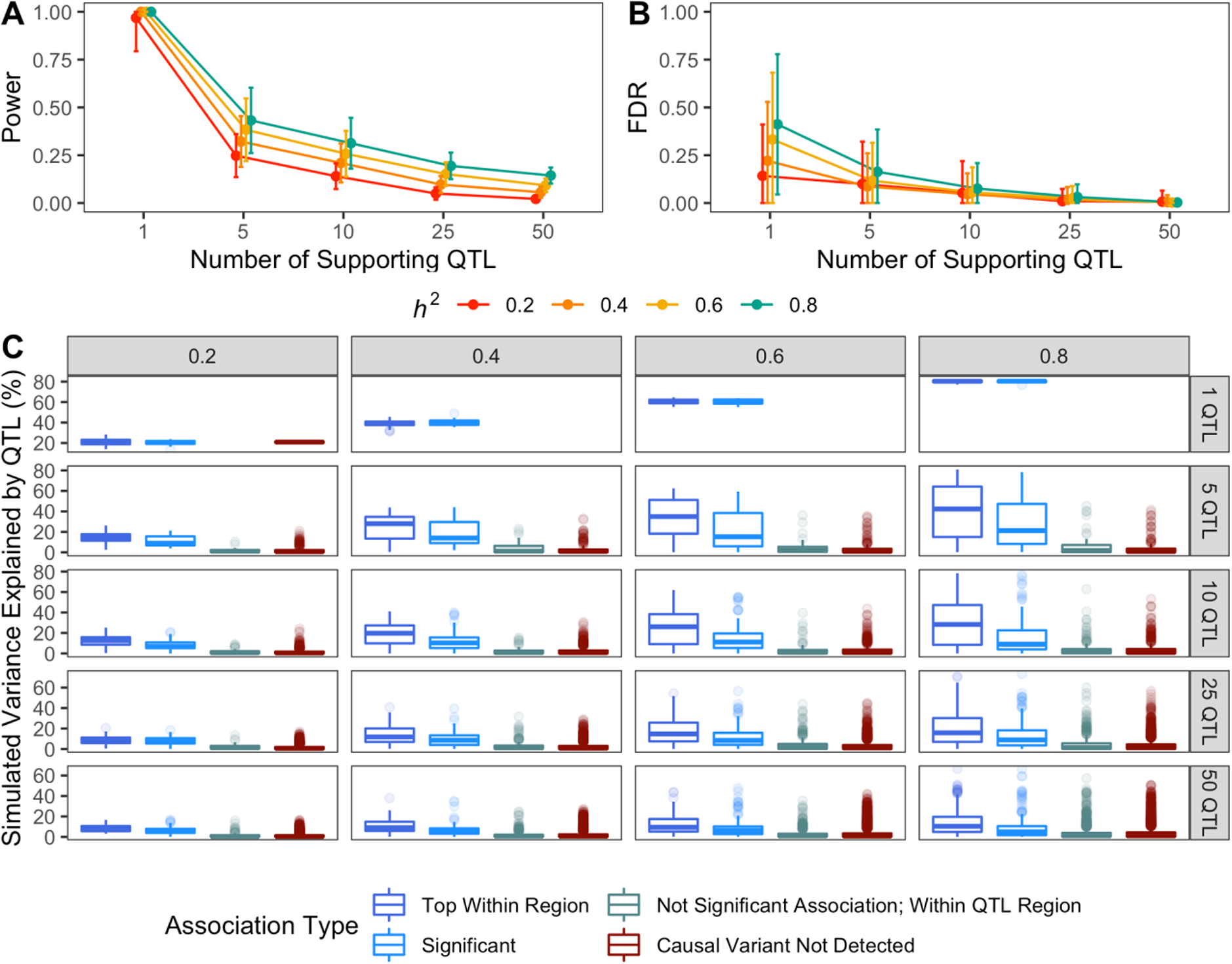
Performance benchmarks for GWA mapping of complex traits in *C. elegans*. Estimates of power (A) and false discovery rate (B) as a function of the narrow-sense heritability [0.2 (red), 0.4, (orange), 0.6 (yellow), 0.8 (green)] and number of causal QTL (ranging from 1-50 QTL) underlying quantitative traits (x-axis). (C) The empirical phenotypic variance explained by each simulated QTL among all architecture regimes, broken out by whether the causal QTL was the top association within a QTL region of interest (dark blue), significant (and thereby exceeding the threshold of significance by multiple testing, light blue), or not a significant association but residing within the QTL region of interest (slate grey) or outside any region of interest (red). Lines stretching from each point represent the standard deviation of the performance estimate among all replicate mappings in (A) and (B). Square boxes linked to black dots in (C) contain the median simulated variance explained by each QTL for that association category within an architecture regime.

**Figure 2:**
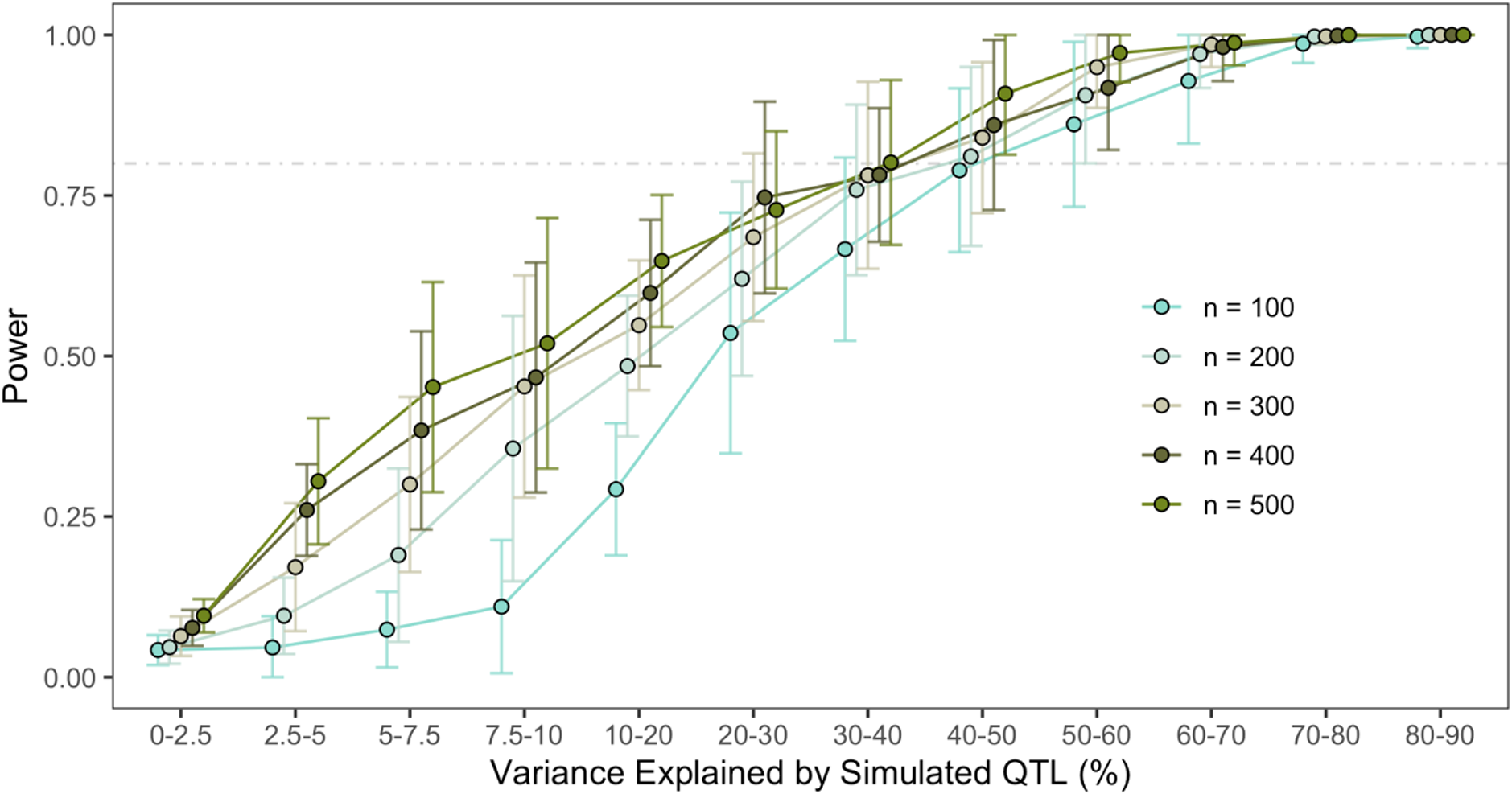
Impact of sample size and strain selection on sensitivity of QTL detection. Power estimates (A) for GWA mappings conditioning on the variance explained by underlying QTL as a function of sample size and strain selection are shown. The corresponding breakdown of the abundance of QTL explaining increasing phenotypic variance (B) and the minor allele frequencies (MAF, C) of these QTL are shown.

**Table 1:**
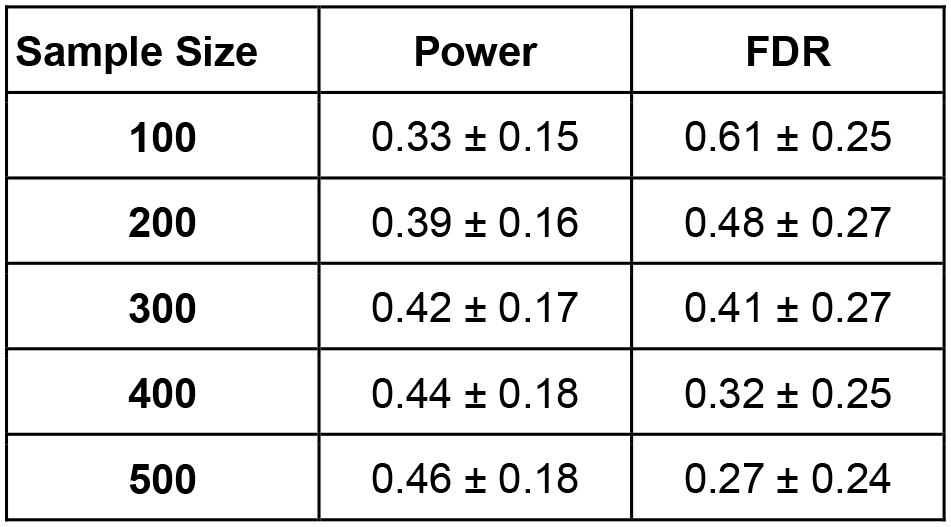
Power and FDR estimates for GWA mappings performed with subsampled populations of increasing depth.

We then measured GWA mapping performance in sets of strains that were distinguished by presence of haplotypes shaped by past selective sweeps (Andersen *et al*. 2012; Crombie *et al*. 2019; Zhang *et al*. 2021). Using the criterion of whether strains harbored at least one chromosome composed of at least 30% swept haplotypes, we divided the 540 strains into two groups: “swept” strains (*n* = 357) and “divergent” strains (*n* = 183). We then simulated and mapped 50 quantitative traits supported by 10 QTL and *h^2^* = 0.8, and QTL effects were once again sampled from a *Gamma* (*k =* 0.4, *θ* = 1.66) distribution. We performed these simulations using populations of equal sampling depth (*n =* 144) from swept strains, divergent strains, and 144 randomly sampled strains from the entire CeNDR strain collection.

We observed that strain selection has a large impact on the sensitivity with which QTL of varying importance are detected. We also observed that the power to detect QTL explaining increasing amounts of phenotypic variance differed dramatically between mappings among strains with similar genome-wide signatures of positive selection and randomly subsampled populations of equal depth (**Figure 3A**). Two patterns emerged from these results. First, swept populations exhibited greater detection power than other populations for QTL that explained greater than 10% of the phenotypic variance. Furthermore, for QTL that explained more than 20% of the phenotypic variance, swept strains exhibited roughly 95% power and other populations exhibited less than 62% power (**Supplemental Table 5**). Second, for QTL explaining greater than 20% of the phenotypic variance, populations assembled without regard for selective sweep haplotypes exhibited lower power than both swept and divergent populations, despite divergent populations having, on average, lower minor allele frequencies of detected and simulated QTL with detected QTL explaining similar amounts of phenotypic variance (**Figure 3B,C**). Nevertheless, these initial simulated mappings provide evidence that strain choice as well as sampling depth dictate the realized genetic architecture of *C. elegans* quantitative traits.

**Figure 3:**
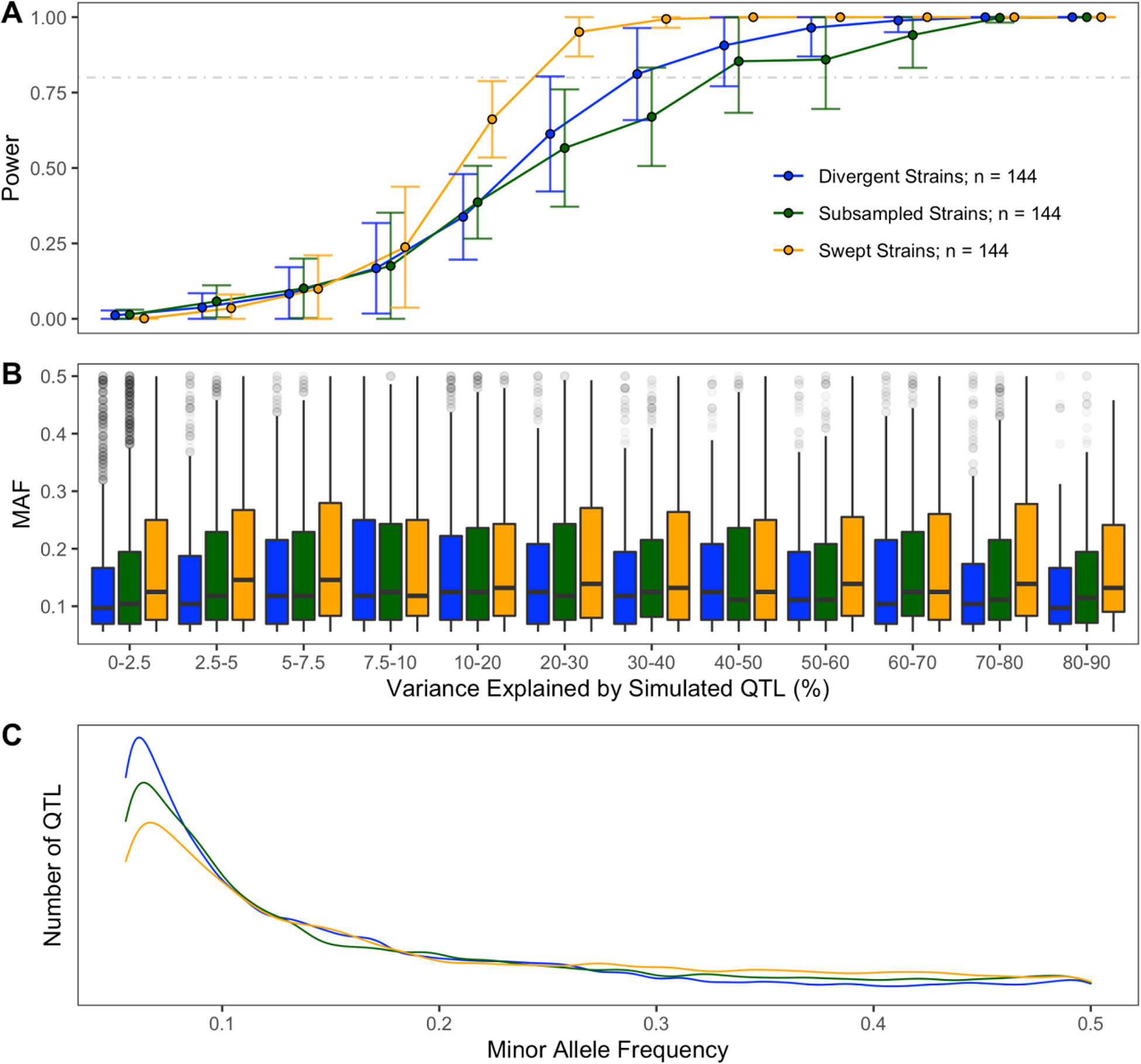
Population composition alters performance and underlying distribution of variants. The fraction of simulated QTL detected by GWA (A) and their minor allele frequencies (B) are plotted as a function of the variance they explain and strain selection. (C) The underlying distributions of minor allele frequencies and effects of all simulated QTL for each population are displayed.

### Fine-scale genomic landscape of GWA performance in C. elegans

The genomes of *C. elegans* wild isolates have been heavily shaped by the evolution of self-fertilization. The recombination rate across the arms of chromosomes is significantly higher than across centers (Rockman and Kruglyak 2009). Many *C. elegans* strains harbor selective sweep haplotypes from which recent adaptation to human-associated niches has purged genetic diversity (Andersen *et al*. 2012; Zhang *et al*. 2021) and hyperdivergent regions that maintain the variation necessary for evolvability (Lee *et al*. 2021). Selective sweep and hyperdivergent region haplotype frequencies and distributions vary across wild isolates, motivating us to ask whether heterogeneity in GWA sensitivity among populations with different demographics can be partly explained by which chromosomes QTL are located and whether these QTL are also located in hyperdivergent regions. In order to assess these points, we simulated 100 mappings of a single QTL with a defined effect size in a population of 182 swept strains and a population of 183 divergent strains. For each set of 100 mappings, the locations of the simulated QTL were constrained to i) a particular chromosome, ii) the region of the chromosome (arms or centers), or iii) within or outside of divergent regions. For each mapping, the heritabilities of the simulated traits were also set to 0.2, 0.5, or 0.8.

We observed several critical differences in mapping performance across different regions of the genome and between divergent and swept mapping populations (**Figure 4A**). At low trait heritability, power to detect QTL was significantly lower among divergent strains than swept strains across all chromosomes, regardless of whether they were in divergent regions, arms, or centers of the chromosome (Kruskall-Wallis test; *p* < 0.0004). We also observed subtle differences in the relative detection power for QTL within certain chromosomes within these classes (**Supplemental Table 6**). Strain sets exhibited identical power to detect QTL genome-wide when *h^2^* = 0.8. The empirical false discovery rate of mappings was significantly greater in mappings among divergent strains than swept strains regardless of the location of simulated QTL (Kruskall-Wallis test; *p* < 0.00001). These differences are likely caused by the large extent to which the divergent population was structured into distinct clusters (**Figure 4B**), and the swept population much closely approximates a star phylogeny because most variation in the population segregates on a much more common genetic background of swept haplotypes (**Figure 4C**). These results confirm a clear effect of population structure and evolutionary history in the species on both genome-wide precision and local detection power of GWA mapping.

**Figure 4:**
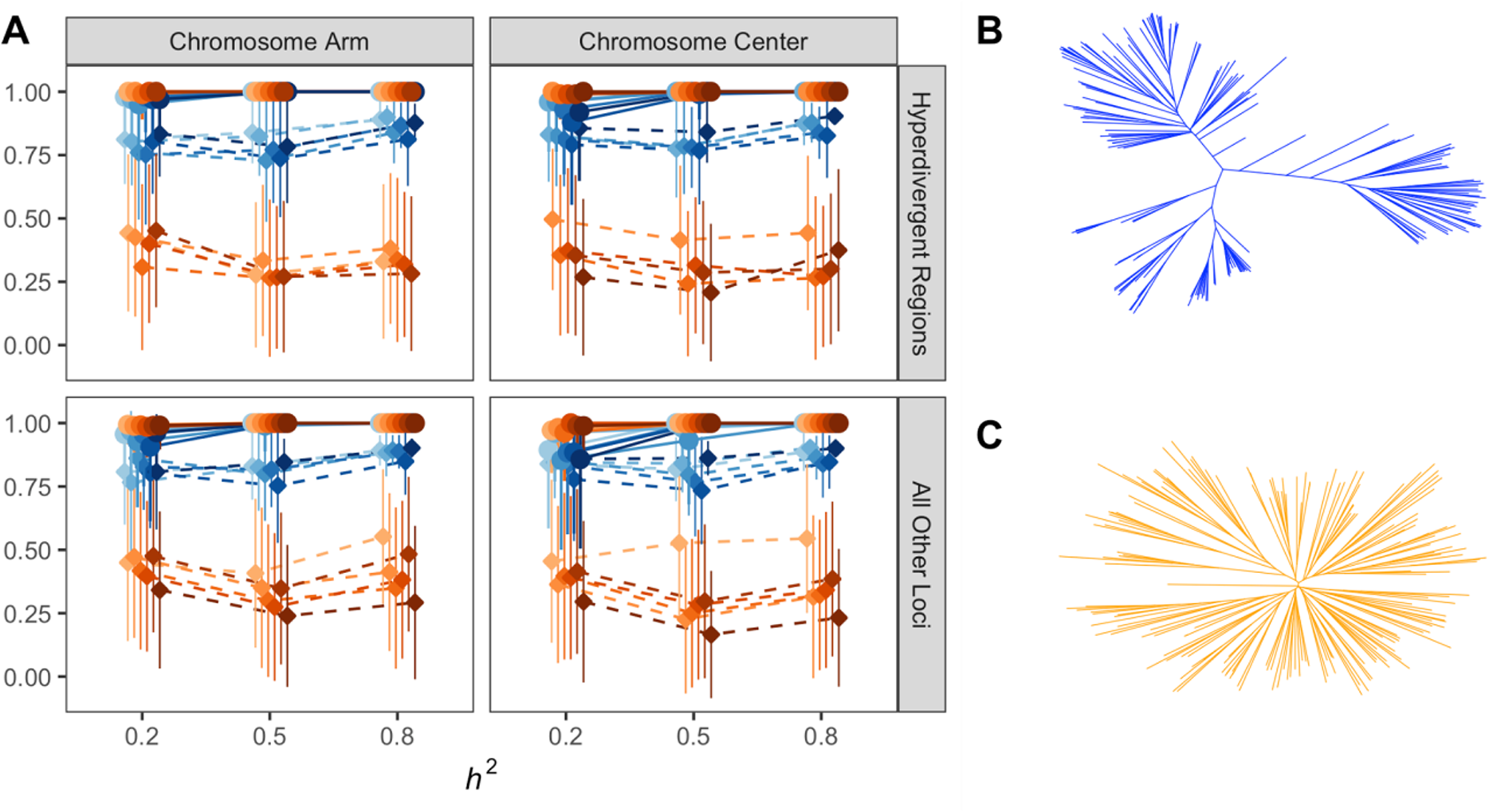
Evolutionary history dictates the fine-scale landscape of GWA performance. A) The mean fraction of simulated QTL detected by GWA (circles, solid lines) and the empirical FDR (diamonds, dashed lines) are plotted as a function of different genomic locations where QTL were simulated: among hyperdivergent regions with respect to the N2 reference genome, or among all other loci, as well as on the low-recombination centers or high-recombination arms of chromosomes. Shading of blue and orange points in A) corresponds to chromosome I (lightest) to chromosome X (darkest) in order. The phylogenetic relationship of each mapping population are shown in B) (183 divergent strains, blue) and C) (182 swept strains, orange).

We also investigated whether certain genomic regions provided varying performance for GWA mapping in *C. elegans*, motivated by the observation of varying population recombination rates on the arms and centers of chromosomes (Rockman and Kruglyak 2009), common selective sweep haplotypes in certain *C. elegans* populations (Andersen *et al*. 2012; Zhang *et al*. 2021), and hyperdivergent haplotypes that segregate among wild strains (Lee *et al*. 2021). Within the swept population, we observed no significant differences in power to detect QTL simulated in hyperdivergent regions nor on chromosome arms compared to centers (*h^2^* = 0.2, Kruskall-Wallis test; *p* = 0.0795). By contrast, power to detect QTL within the divergent population differed as a function of whether they were simulated in hyperdivergent regions or different parts of the chromosome (*h^2^* = [0.2, 0.5]; Kruskall-Wallis test, *p* < 0.0001; Dunn test, *p_adj_* < 0.02) (**Supplemental Table 7**). Once again, the empirical false discovery rate among divergent regions and different chromosomal regions varied significantly for all trait heritabilities within both the divergent and swept strain set (Kruskall-Wallis test; *p* < 0.02) (**Supplemental Table 8**).

Finally, we asked whether GWA mapping performance varied between chromosomes controlling for historic recombination rate differences or the population divergence of haplotypes. We only observed one case where detection power varied significantly among chromosomes - power to detect QTL outside of hyperdivergent regions on the center of chromosome III was significantly lower than that observed for chromosomes I, IV, V, and X at *h^2^*= 0.5 (Dunn test, *p_adj_* ≤ 0.0103) among divergent strains (**Supplemental Table 9**). Notably, this chromosome also harbors the fewest sweep haplotypes in the *C. elegans* population, which could indicate that this local dip in power could be caused by a local enrichment of rare haplotypes among more divergent strains in the population. Empirical FDR varied significantly among chromosomes in several instances among both divergent and swept strain sets (Kruskall-Wallis test; *p* < 0.05) (**Supplemental Table 10**). Taken together, these results demonstrate that differences in GWA mapping performance arising from strain composition differences are likely caused in part by the unique patterns of genetic variation throughout the *C. elegans* genome.

### NemaScan recapitulates previously validated genetic associations

Previous work has used GWA mappings to identify QTL and subsequently identify quantitative trait variants (QTV) in *C. elegans* (Evans et al. 2021b). In order to test whether NemaScan performs similarly in practice to cegwas2-nf, the previous mapping pipeline (https://github.com/AndersenLab/cegwas2-nf) that used the EMMA algorithm (Kang *et al*. 2008) implemented by R/rrBLUP (Endelman 2011), we re-mapped five quantitative traits using both cegwas2-nf and NemaScan. Raw trait files were downloaded from the supplemental materials for each published mapping and re-mapped using the 20210121 CeNDR release VCF. In each case, the major QTL underlying each trait were mapped using both platforms (**Figure 5A**). Of the 16 QTL identified across the previously mapped traits, 14 were recovered by NemaScan. Furthermore, in some instances NemaScan was qualitatively more specific with respect to QTL identification. For example, in the original mapping of arsenic resistance, two QTL in significant LD were identified on chromosomes I and III. Because these sets of markers have identically significant association scores across the interval, the most likely cause of this association is that population structure among the phenotyped strains is causing an entire shared haplotype to be tagged as significant. When mapped with NemaScan, the significance of this association was slightly lower than that of the previous mapping. Similarly, the two previously mapped abamectin resistance QTL were detected and assigned greater significance by NemaScan (**Supplemental Figure 5**). These findings confirm that NemaScan has sufficient detection power to recapture known genetic architectures of real traits, including many with empirically proven QTV. Among each of these mappings, we observed that the aggregated NemaScan *p-*values (the collection of top associations from either the LMM-EXACT-INBRED or the LMM-EXACT-LOCO algorithm for each marker) exhibited varying levels of inflation relative to both the EMMA mappings and to expected *p-*values for each trait (**Figure 5B**). Although the relative inflation of arsenic resistance, etoposide resistance, and telomere length association mapping statistics were relatively similar, mappings of abamectin resistance and dauer pheromone responses were quite different. This difference can be ascribed in part to the fact that mapping statistics derived from NemaScan are the maximum between two matrix construction options and that, when we compared each set of algorithm-specific raw *p*-values to their expected quantiles, one of the algorithms often displayed less inflation. However, in some cases, like abamectin resistance, the algorithm producing lower *p-*values failed to detect any significant QTL (**Supplemental Figure 6**), indicating that the flexibility of algorithm choice in NemaScan mappings could be a source of strength when population structure of phenotypes interacts with trait heritability to have an outsized influence on QTL detection.

**Figure 5:**
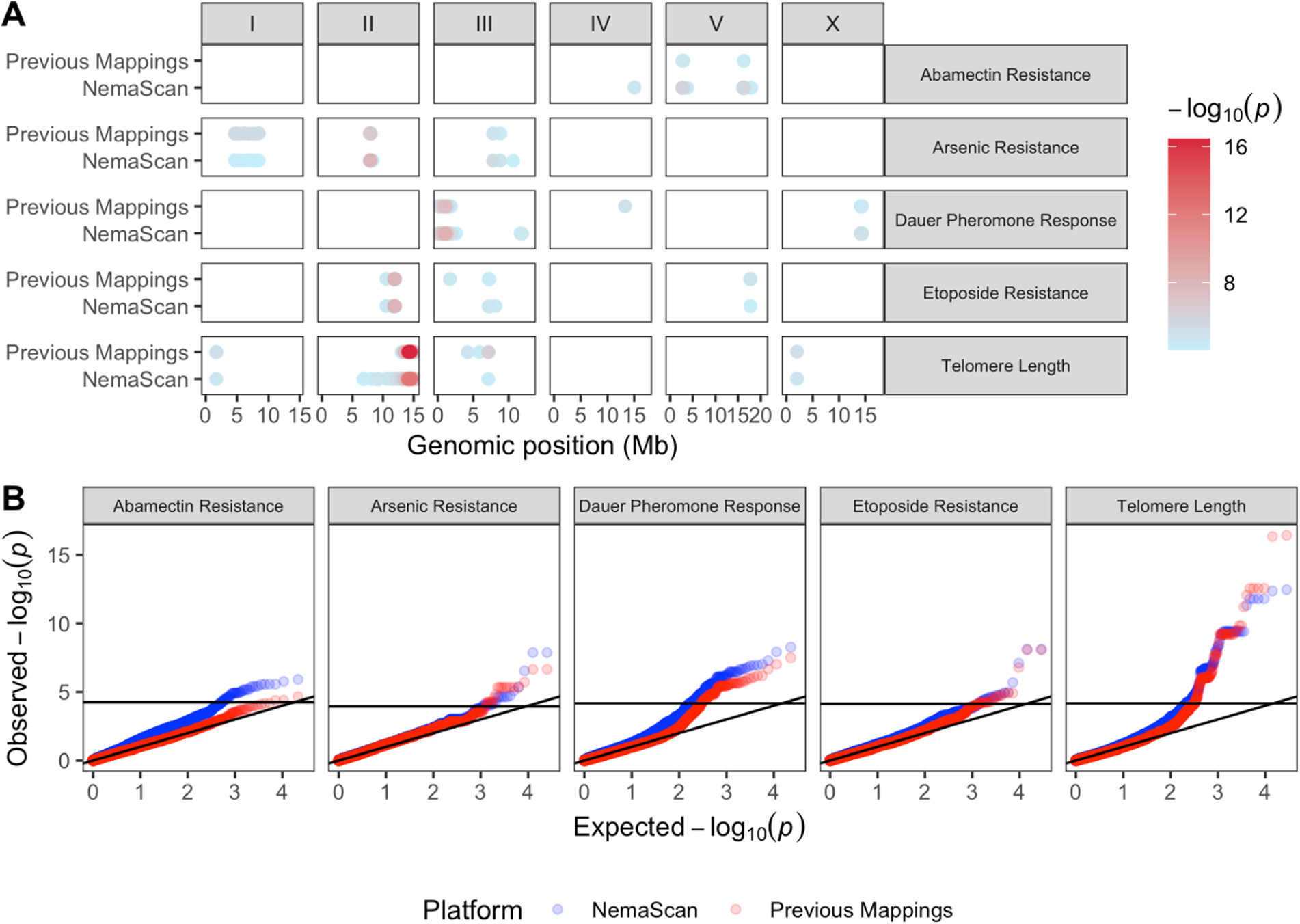
GWA mapping with NemaScan recaptures previously validated QTVs. A) Significant genetic associations are shown genome-wide for five quantitative traits that were re-mapped using the 20210121 CeNDR release both with cegwas2-nf (“Previous Mappings”) and NemaScan, and the strength of the association is displayed increasing from blue to red. B) Quantile-quantile plots of all -log transformed *p*-values are plotted against their expected rank, with the horizontal line in each panel indicating the trait-specific multiple testing correction significance threshold.

## DISCUSSION

### GWA mapping as a tool for QTL discovery in C. elegans

The *C. elegans* community has contributed steadily to the catalog of species-wide genetic variation. As the number of genetically characterized unique strains expands the CeNDR collection, we learn more about genomic patterns of diversity all over the world. The prospects for using GWA mapping to dissect the genetic underpinnings of complex traits have improved in tandem. Although the community has successfully employed GWA mappings in *C. elegans* to discover novel genes related to a variety of traits, we lack a robust characterization of the power and precision with which this resource is equipped to detect QTL. Evaluating population-based genetic resources for other systems using simulations has provided key benchmarks for their respective communities (Kover *et al*. 2009; Bennett *et al*. 2010; King *et al*. 2012a; b; Bouchet *et al*. 2017; Noble *et al*. 2017; Gage *et al*. 2018; Keele *et al*. 2019). The burgeoning *C. elegans* quantitative genetics community has applied GWA mapping to a growing collection of wild strains and identified genetic variants linked to complex traits with novel biomedical and evolutionary implications. In the simulations presented here, we systematically tested a robust framework for GWA against a variety of genetic architectures and sample populations to contextualize past, present, and future studies using CeNDR. However, some important limitations of our simulation framework have implications in real populations. First, simulated causal variants were selected from the minor allele frequency and LD-filtered variant set, meaning that all QTL are perfectly tagged and at greater than 5% frequency in the population, upwardly biasing their detection in simulations. In practice, GWAS may underestimate the effects of rare QTVs imperfectly tagged by filtered variants or fail to detect these variants altogether. Future work should prioritize rare variant detection, especially given their implied frequency in divergent populations (**Figure 3C**). Second, effects assigned to simulated causal variants were drawn from a *Gamma* (*k =* 0.4, *θ* = 1.66) distribution (**Supplementary Figure 3**) creating genetic architectures heavily biased against detection of causal alleles with very small effects. In practice, traits supported by fewer QTL of greater effect will be more amenable to GWA mapping, even at low heritability (**Figure 1C**). In spite of these limitations, we hope to provide the community with a flexible platform for QTL detection and simulation-based performance evaluation.

Similar to multiparent mapping populations in other systems, we confirmed that the prospects of identifying QTL that explain a less than substantial proportion (∼10%) of overall trait variance depend primarily on three factors: (1) the number of strains being phenotyped, (2) the precision with which phenotypes can be measured, and (3) the composition of the mapping population. For instance, we observed that measuring only 100 wild isolates is expected to provide almost 80% power to detect QTL that explain greater than 40% of the phenotypic variance. For many traits, it is no small feat to measure 100 strains with sufficient replication for line means to robustly represent that genetic background in a GWA mapping population. A recent GWA analysis of sperm size among 96 wild strains and N2 revealed no significant associations despite the nomination of the candidate gene *nurf-1* using segregating mutations between the N2 and LSJ lineages (Gimond *et al*. 2019). Another recent GWA analysis of starvation resistance using population RAD-seq read abundance in a 96 strain co-culture revealed a single large-effect QTL on chromosome III whose effect was validated using near-isogenic lines and was present in 11% of wild strains (Webster *et al*. 2019). These applications of GWA mappings represent mixed outcomes, providing some practical support for the conclusions of our simulations – lower sampling depths are not expected to capture entire genetic architectures, including small-effect loci or impactful alleles that segregate at low frequency (less than 5% of the population). Larger sample sizes (300-500 strains) and potentially less experimentally strenuous trait measurements are optimal for identifying loci that confer more modest effects (roughly 5-10% of the phenotypic variance) with greater likelihoods. Traits that can be measured in high-throughput (Hahnel *et al*. 2018; Evans *et al*. 2021a) or as intermediate traits (*e.g.*, mRNA abundances) lend themselves to dissection in hundreds of strains and QTL conferring more subtle effects can be more easily resolved. At the current size of CeNDR, the primary driver of sampling depth of GWA mapping populations should be the balance between phenotyping effort for the trait of interest and the end goal of association mapping given the roughly estimated heritability of the trait (**Figure 2**) and the lower bound of the effect of QTL that will be detected (**Figure 3**). In many cases, evaluating the same trait using linkage mapping in complementary populations (*i.e.*, traits segregate similarly between parental strains of the cross and in the association mapping population) can validate effect sizes and provide additional support for candidates from GWA (Zdraljevic *et al*. 2019; Webster *et al*. 2019; Evans *et al*. 2021a).

### Population structure is a major determinant of performance

In this study, we also quantified the impact of mapping population structure on the power and precision of GWA mapping. In comparing mappings derived from (1) choosing strains from CeNDR at random, (2) swept strains, and (3) divergent strains of equal sampling depth, we confirmed that the most power to map QTL was provided by sampling swept strains (**Figure 3A**). We also found from these comparisons that the empirical FDR among the divergent strain mappings was significantly higher than the swept strain mappings when a single QTL was simulated (**Figure 4A**). This result aligns with outcomes of past GWA analyses in model organisms, wherein mappings among structured populations provided less specific inference of genetic architectures (Kang *et al*. 2008). *C. elegans* populations also harbor highly variable patterns of genetic variation across the genome in these distinct populations, which contribute to subtle differences in local performance and inference of associations (**Figure 4A**). However, we chose only one collection of strains to represent both divergent and swept mapping populations when considering local performance differences, which limits the general extensibility of these particular benchmarks in other populations. As different combinations of strains with varying landscapes of selective sweeps and hyperdivergent regions are tested, we will learn more about the relative influences of these regions on performance. Before concluding that an experimenter’s particular mapping population will be less powerful because it contains many divergent strains, one is advised to perform their own population-specific simulations. Below, we outline some limitations to pursuing GWA in only swept strains in certain contexts.

First, trait heritability is a major driver of detection power, which means that if the phenotype of interest does not vary significantly among swept strains, the prospects for mapping its genetic architecture heavily rely on low experimental noise. Divergent strains have been shown to exhibit distinct population-wide phenotypic differences from swept strains (Zhang *et al*. 2021) and therefore might be expected to contribute significantly to estimates of narrow-sense heritability of other traits. Second, swept populations will be enriched for alleles that have arisen relatively recently on swept haplotypes. Some QTL will be slightly more common in the population in swept populations (**Figure 3C**), but swept populations provide a limited view of whether these QTL identified are meaningful in divergent populations that are more representative of the ancestral niche of *C. elegans* (Lee *et al*. 2019, 2021; Crombie *et al*. 2019). We know of many examples where strains more closely associated with human colonization and laboratory domestication express trait differences uncharacteristic of “wild” *C. elegans* isolates (Sterken *et al*. 2015; Schulenburg and Félix 2017). Third, one kinship matrix construction algorithm used in our GWA platform was designed, in part, to collapse extremely close relatedness among inbred individuals by creating sparse genetic covariance. This calculation is expected to provide more power in swept populations than divergent populations because the covariance among swept strains will be small enough for the algorithm to collapse more often than among divergent strains.

A helpful comparison for the prospects of *C. elegans* GWAS is the successes of identifying disease risk alleles in human populations. *Trans*-ethnic GWAS has successfully identified common variants linked to complex human diseases by leveraging rich data and population sizes (Wojcik *et al*. 2019; Pendergrass *et al*. 2019; Hu *et al*. 2021). However, generalized predictions of disease risk in the form of polygenic risk scores suffer from sampling bias, genetic heterogeneity, and varying frequencies of risk alleles among distinct subpopulations (Li and Keating 2014; Márquez-Luna *et al*. 2017; Martin *et al*. 2019, 2020). As the community sampling of diverse *C. elegans* strains grows, GWAS will provide more power to detect QTL with more modest effects, and we will achieve more success in identifying common genetic variants linked to complex traits. However, one advantage of *C. elegans* is that complementary techniques for quantitative genetics are easily achievable and essential for validating candidate loci from GWA mappings. Near-isogenic lines (NILs) and recombinant inbred lines (RILs) can be derived from individual strains with large phenotypic contrasts and used for fine mapping alleles, making hypothesis-driven inferences of GWA candidate gene identification and functional tests more addressable than could be hoped for in many other species. As genomic resources for comparative evolutionary studies in *C. elegans* grow, we will characterize hyperdivergent regions more completely so that variants identified in GWA within these regions can be more confidently nominated as candidates. Furthermore, future endeavors of GWA mapping should explicitly control for the extensive population structure present among divergent strains using statistical techniques being actively applied to significantly larger cohorts of stratified human populations (Wojcik *et al*. 2019).

## ACKNOWLEDGEMENTS

We would like to thank members of the Andersen laboratory and Dr. Matthew Rockman for helpful comments on the manuscript. This work was supported by a Research Grant to E.C.A. from HFSP (Ref.-No: RPG0001/2019).

## Supplemental Figures

**Supplemental Figure 1:**
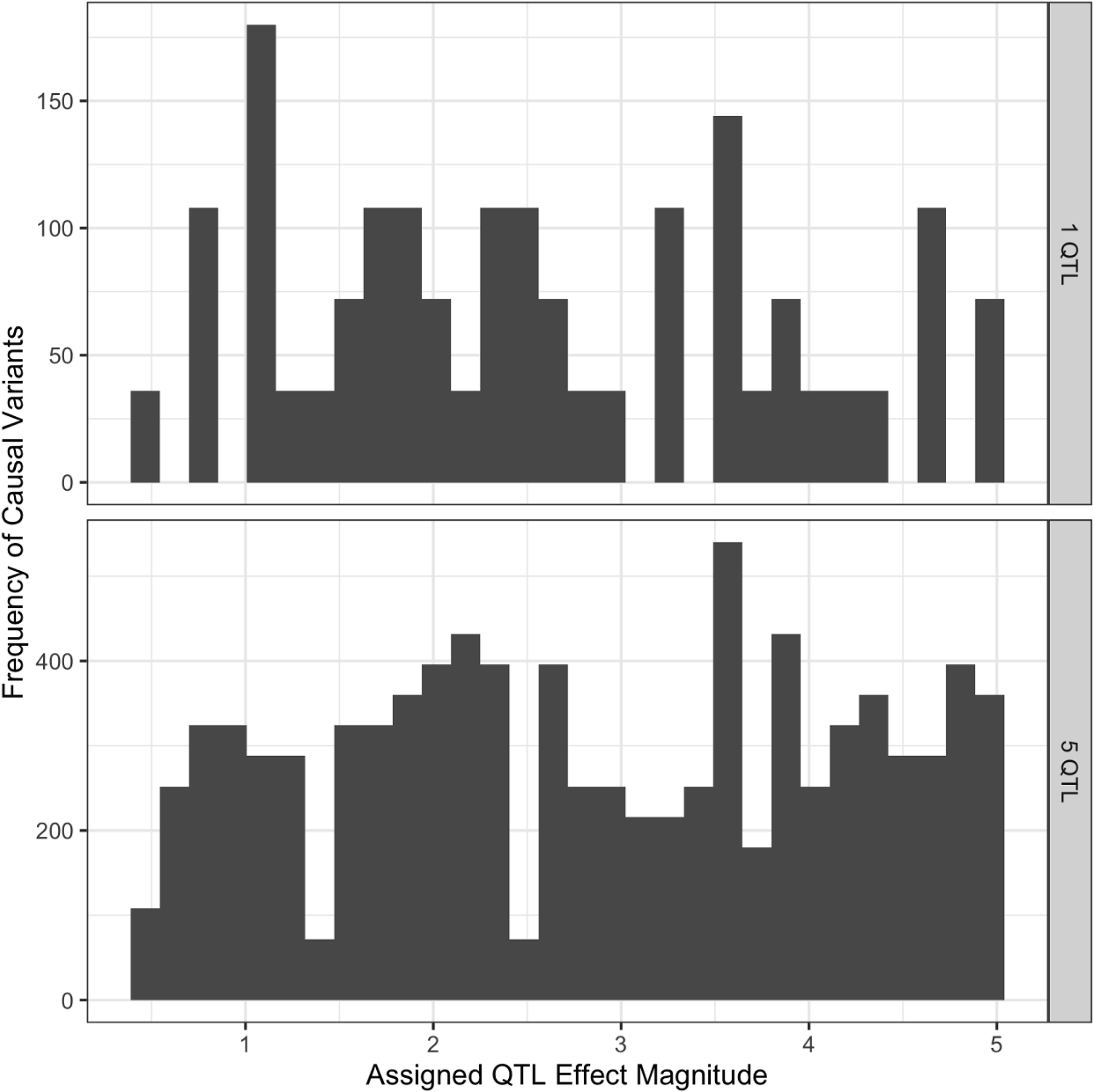
Effect size distribution of simulations comparing algorithm performance

**Supplemental Figure 2:**
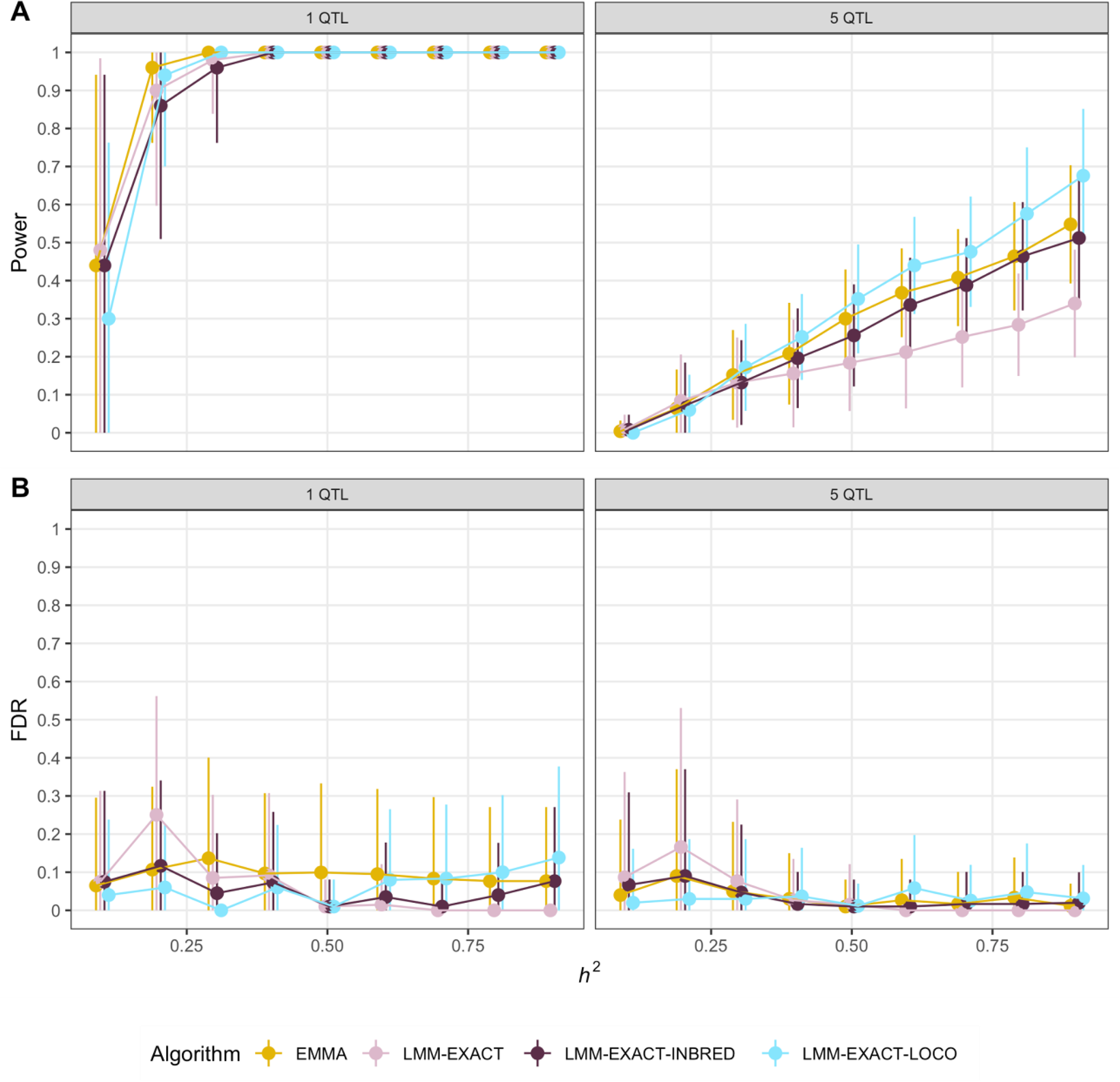
Power and false discovery rate of GWA mapping across various algorithms

**Supplemental Figure 3:**
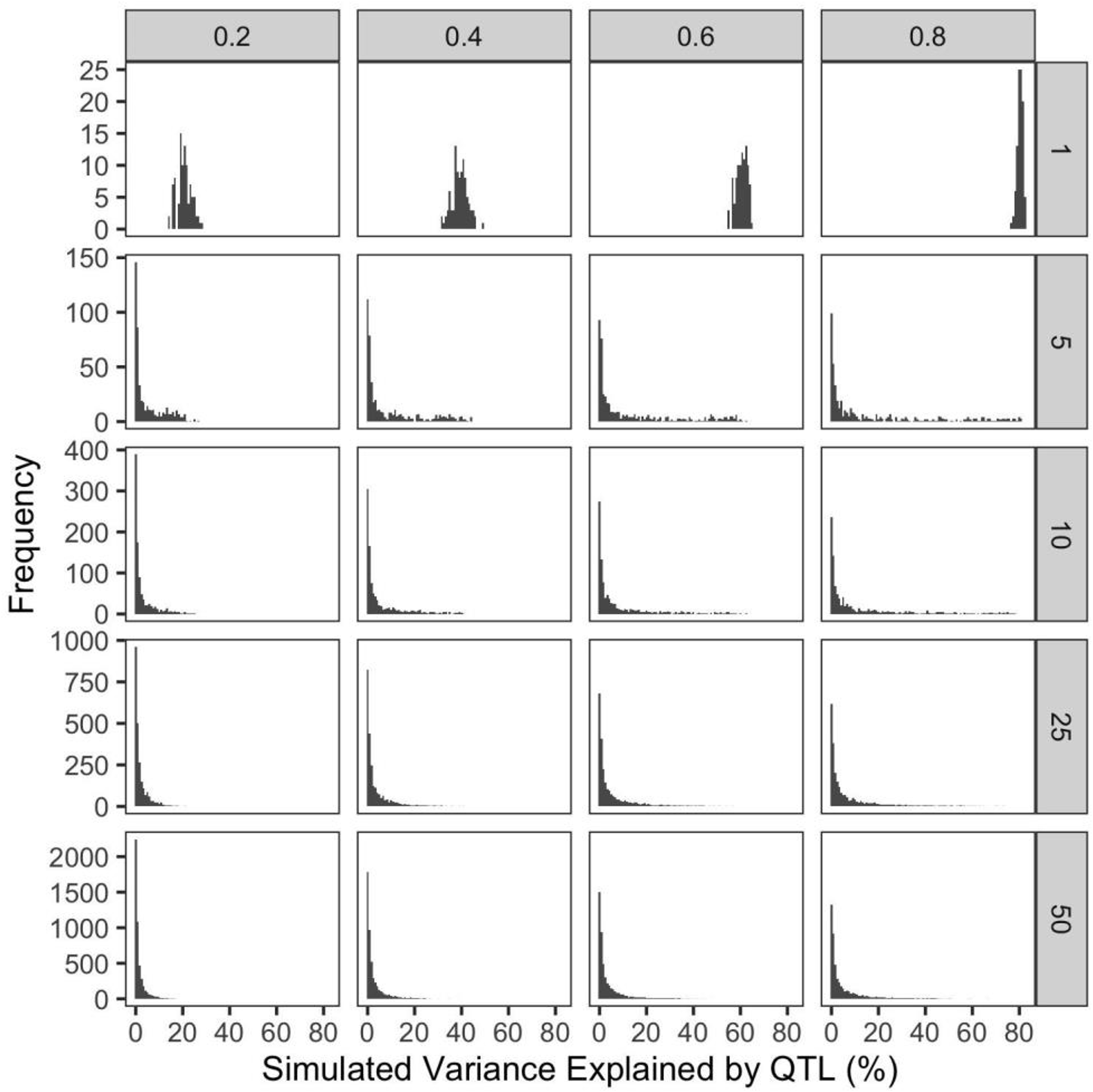
Distributions of simulated QTL effects expressed as the fraction of phenotypic variance explained. Horizontal panels denote the number of simulated QTL per trait and vertical panels denote the heritability of each simulated trait.

**Supplemental Figure 4:**
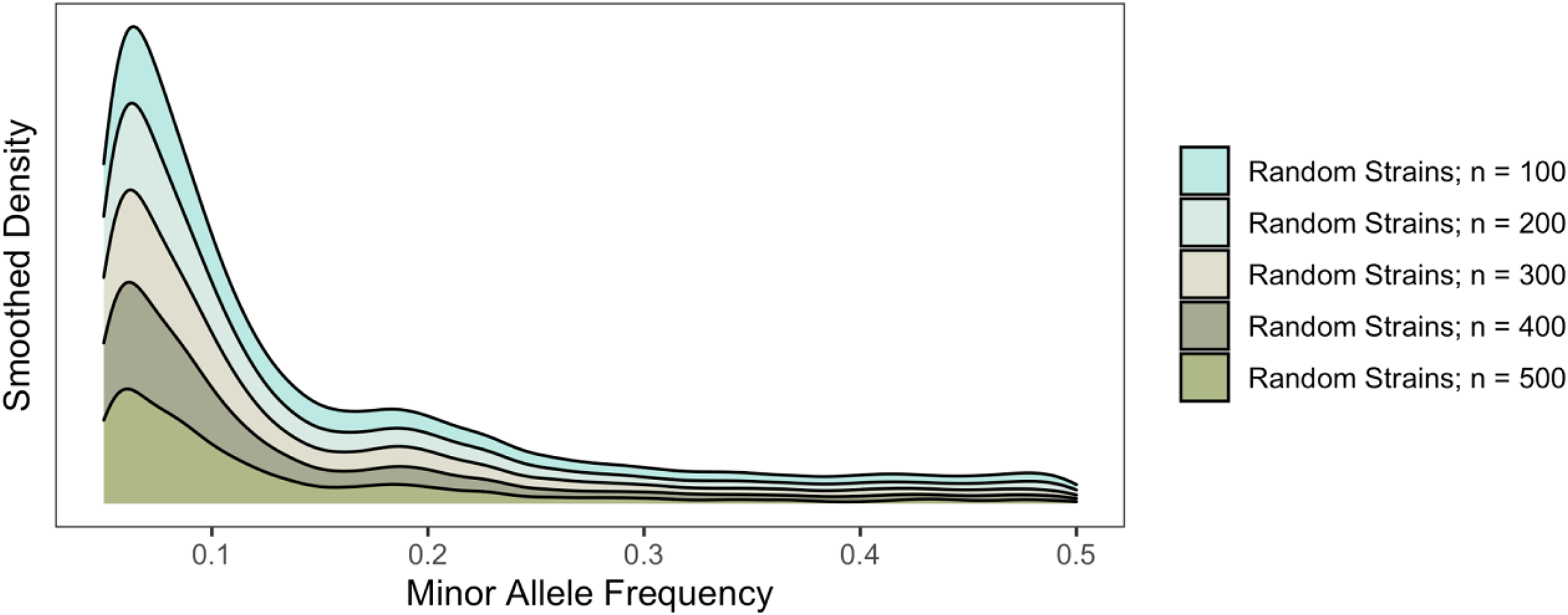
Distributions of all simulated QTL minor allele frequencies among mapping populations of increasing size

**Supplemental Figure 5:**
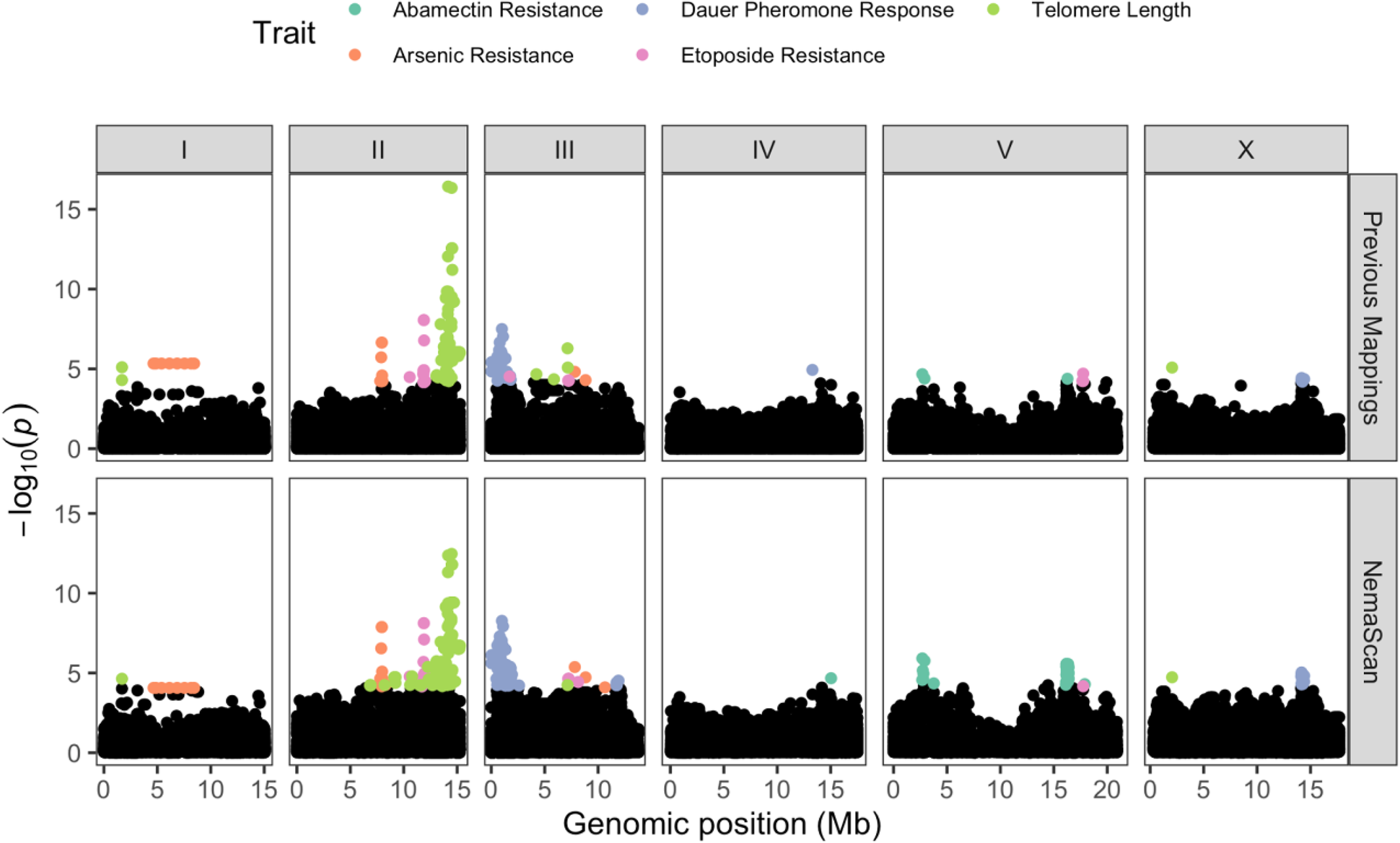
Manhattan plots of previous GWA mappings and NemaScan mappings. Markers exceeding the multiple testing correction threshold are colored according to the mapped trait of interest.

**Supplemental Figure 6:**
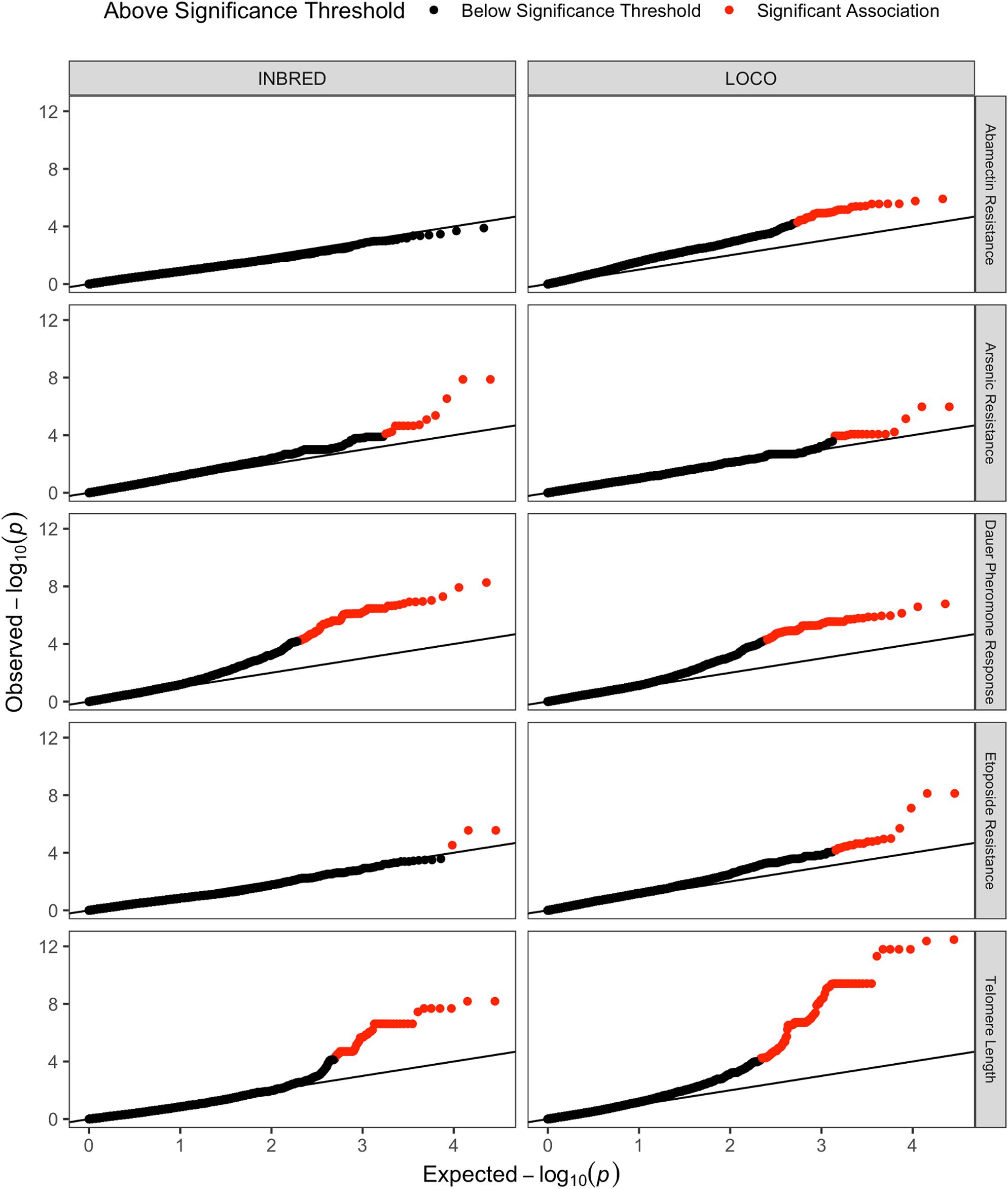
QQ plots of raw NemaScan GWA mappings corresponding to the mapping algorithm that generated association scores for each trait, colored by whether the significance of each association exceeds the multiple testing threshold.

## Supplemental Tables

Supplemental Table 1: Simulation summary

Supplemental Table 2: Differences in power to detect QTL between mapping algorithms at increasing heritability for one and five underlying QTL.

Supplemental Table 3: Differences in empirical FDR between mapping algorithms at increasing heritability for one and five underlying QTL.

Supplemental Table 4: Average power to detect QTL explaining increasing phenotypic variance among subsampled populations of increasing sampling depth

Supplemental Table 5: Average power to detect QTL explaining increasing phenotypic variance among 144 randomly sampled divergent strains, 144 randomly sampled swept strains, and 144 randomly sampled strains from the overall CeNDR population

Supplemental Table 6: Differences in power to detect QTL between different chromosomes controlling for hyper-divergence and historic recombination groups (arms vs. centers)

Supplemental Table 7: Power to detect simulated in hyperdivergent regions or different parts of the chromosome within the mapping populations

Supplemental Table 8: Empirical FDR of mappings as a function of whether QTL were simulated in divergent regions and different chromosomal regions

Supplemental Table 9: Power to detect simulated QTL on different chromosomes, within hyperdivergent regions, historic recombination groups, and strain sets

Supplemental Table 10: Empirical FDR of mappings as a function of whether QTL were simulated on different chromosomes, within hyperdivergent regions, historic recombination groups, and strain sets

